# Genome-wide association study of Parkinson’s disease progression biomarkers in 12 longitudinal patients’ cohorts

**DOI:** 10.1101/585836

**Authors:** Hirotaka Iwaki, Cornelis Blauwendraat, Hampton L. Leonard, Jonggeol J. Kim, Ganqiang Liu, Jodi Maple-Grødem, Jean-Christophe Corvol, Lasse Pihlstrøm, Marlies van Nimwegen, Samantha J. Hutten, H. Nguyen Khanh-Dung, Jacqueline Rick, Shirley Eberly, Faraz Faghri, Peggy Auinger, Kirsten M. Scott, Ruwani Wijeyekoon, Vivianna M. Van Deerlin, Dena G. Hernandez, J. Raphael Gibbs, Kumaraswamy Naidu Chitrala, Aaron G. Day-Williams, Alexis Brice, Guido Alves, Alastair J. Noyce, Ole-Bjørn Tysnes, Jonathan R. Evans, David P. Breen, Karol Estrada, Claire E. Wegel, Fabrice Danjou, David K. Simon, Ole Andreassen, Bernard Ravina, Mathias Toft, Peter Heutink, Bastiaan R. Bloem, Daniel Weintraub, Roger A. Barker, Caroline H. Williams-Gray, Bart P. van de Warrenburg, Jacobus J. Van Hilten, Clemens R. Scherzer, Andrew B. Singleton, Mike A. Nalls

## Abstract

**Background:** Several reports have identified different patterns of Parkinson’s disease progression in individuals carrying missense variants in the *GBA* or *LRRK2* genes. The overall contribution of genetic factors to the severity and progression of Parkinson’s disease, however, has not been well studied.

**Objectives:** To test the association between genetic variants and the clinical features and progression of Parkinson’s disease on a genome-wide scale.

**Methods:** We accumulated individual data from 12 longitudinal cohorts in a total of 4,093 patients with 25,254 observations over a median of 3.81 years. Genome-wide associations were evaluated for 25 cross-sectional and longitudinal phenotypes. Specific variants of interest, including 90 recently-identified disease risk variants, were also investigated for the associations with these phenotypes.

**Results:** Two variants were genome-wide significant. Rs382940(T>A), within the intron of *SLC44A1*, was associated with reaching Hoehn and Yahr stage 3 or higher faster (HR 2.04 [1.58, 2.62], P-value = 3.46E-8). Rs61863020(G>A), an intergenic variant and eQTL for *ADRA2A*, was associated with a lower prevalence of insomnia at baseline (OR 0.63 [0,52, 0.75], P-value = 4.74E-8). In the targeted analysis, we found nine associations between known Parkinson’s risk variants and more severe motor/cognitive symptoms. Also, we replicated previous reports of *GBA* coding variants (rs2230288: p.E365K, rs75548401: p.T408M) being associated with greater motor and cognitive decline over time, and *APOE* E4 tagging variant (rs429358) being associated with greater cognitive deficits in patients.

**Conclusions:** We identified novel genetic factors associated with heterogeneity of progression in Parkinson’s disease. The results provide new insights into the pathogenesis of Parkinson’s disease as well as patient stratification for clinical trials.

## Introduction

Parkinson’s disease (PD) is clinically defined by its motor features of rigidity, bradykinesia, gait disturbance, and tremor. Although these prominent features are important for diagnosis, patients with PD also suffer from many non-motor features such as constipation, urinary incontinence, orthostatic hypotension, REM sleep behavior disorder (RBD), apathy, hyposmia, and cognitive impairment (Postuma *et al.*, 2015). Moreover, patients develop motor complications, including wearing off and dyskinesia, as side effects of medication. The onset, intensity and progression of these different PD clinical features vary among individuals, and the mechanisms underlying this heterogeneity are not well understood.

Recent genome-wide studies have identified 90 common variants associated with the risk of PD, with an overall heritability estimated to be between 22-27% (Keller *et al.*, 2012; Nalls *et al.*, 2019). While previous studies have indicated the importance of genetic contributions to disease risk, the contribution of genetic factors to PD progression and heterogeneity has not been well studied. Investigating genetic factors associated with disease progression and heterogeneity in disease presentation is an important step in elucidating the underlying molecular mechanisms and identifying better patient stratification in clinical trials (Leonard *et al.*, 2018).

Longitudinal patient cohorts are powerful resources that can be used to explore the impact of genetics on the trajectory of PD-related phenotypes; the inherent precision of repeated measurements over time provides more power to detect these associations. However, the available number of participants in each study is usually not enough to conduct a genome-wide association study (GWAS). In this study, we accumulated 25,254 follow-up visits from 4,093 patients across 12 cohorts (Table 1) and performed meta-analyses of longitudinal GWAS on the progression markers of Parkinson’s disease. Using the results from this meta-analysis, we evaluated how known risk variants, including the 90 recently identified variants for PD (Nalls *et al.*, 2019), *GBA* protein coding mutations, and *APOE* tagging variants were associated with the progression of phenotypes. To maximize the utility of this work to other researchers, we have made all results from this study publicly searchable and available for download. (https://pdgenetics.shinyapps.io/pdprogmetagwasbrowser/)

**Table 1.**
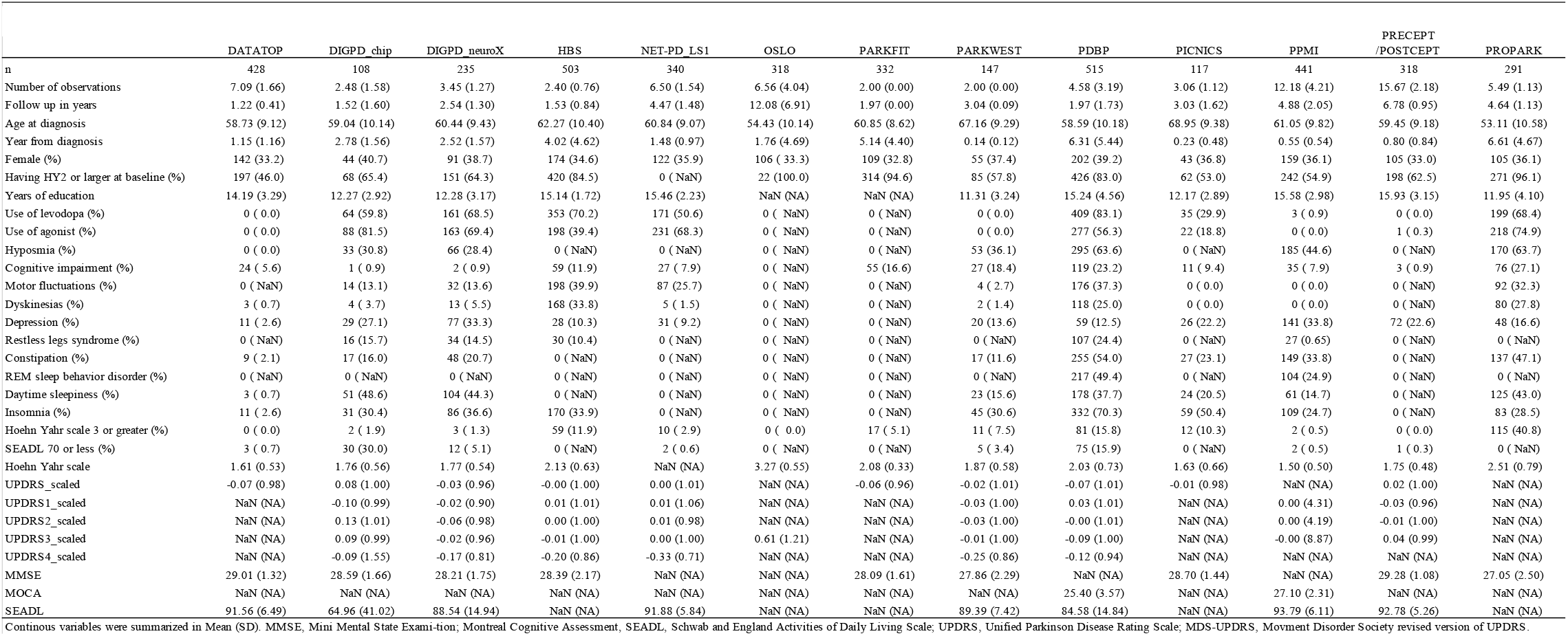
Summary of 13 datasets (12 cohorts)

## Methods

### Cohorts

Twelve longitudinal cohorts of PD patients recruited across North America, Europe and Australia were included in our study. The following observational studies were included: the Drug Interaction with Genes in Parkinson’s Disease (DIGPD), the Harvard Biomarkers Study (HBS), the Oslo Parkinson’s Disease study (partly including retrospective data), the Norwegian ParkWest study (PARKWEST), the Parkinson’s Disease Biomarker Program (PDBP), the Parkinsonism Incidence and Cognitive and Non-motor heterogeneity In Cambridgeshire (PICNICS), the Parkinson’s Progression Markers Initiative (PPMI), and the Profiling Parkinson’s disease study (PROPARK). The four cohorts included were randomized clinical trials: the Deprenyl and Tocopherol Antioxidative Therapy of Parkinsonism (DATATOP), the NIH Exploratory Trials in Parkinson’s Disease Large Simple Study 1 (NET-PD_LS1), the ParkFit study (PARKFIT), and the Parkinson Research Examination of CEP-1347 Trial study with its subsequent prospective study (PreCEPT/PostCEPT). More details of these cohorts are described in Appendix. Participants’ information and genetic samples were obtained under appropriate written consent and with local institutional and ethical approvals.

### Phenotyping

For continuous outcomes, we collected the scores of Hoehn and Yahr staging scale (HY) (Goetz *et al.*,2004), total and sub-scores of the Unified Parkinson’s Disease Rating Scale (UPDRS) or the Movement Disorder Society revised UPDRS version (MDS-UPDRS) (Goetz *et al.*, 2007), Mini-Mental State Examination (MMSE), Montreal Cognitive Assessment (MoCA) (Nasreddine *et al.*, 2005) and the modified Schwab and England Activities of Daily Living Scale (SEADL). With the exception of the subscores of UPDRS/MDS-UPDRS part 4, total scores and the subscores of UPDRS and MDS-UPDRS were normalized to the population-baseline mean and standard deviation and converted to Z values. The subscores of UPDRS/MDS-UPDRS part 4, measuring complication of treatment, were normalized to the mean and standard deviation of all observations because the score was 0 at the baseline for the de-novo PD cohorts. We also determined whether subjects were recorded as presenting the following binomial outcomes during participant visits: constipation, cognitive impairment, depression, daytime sleepiness, Hoehn and Yahr stage of 3 or worse (HY3), hyposmia, insomnia, motor fluctuation, REM sleep behavior disorder (RBD), restless legs syndrome (RLS), and an SEADL of 70 or less (SEADL70). Because study-specific criteria for these binomial outcomes were not consistent amongst the studies, we tried to use the common criteria for these binomial outcomes if we had access to the raw data from the studies. The details of the definitions of binomial outcomes are provided in the Supplemental Table 1.

### Genetics data

The genotyping was conducted with NeuroX, a targeted chip for neurodegenerative disease (Nalls *et al.*, 2015), for NET-PD_LS1, a part of DIGPD (DIGPD_neuroX), HBS, PDBP, and PRECEPT. The rest of DIGPD (DIGPD_chip) were genotyped using Illumina Multi-Ethnic Genotyping Array. Participants in DATATOP, OSLO, PARKFIT, PARKWEST, PICNICS, and PROPARK were genotyped using Illumina Infinium OmniExpress array. Whole genome sequencing data was used for PPMI, with the detailed methods for genome sequencing provided on the PPMI website (https://www.ppmi-info.org/).

Variant inclusion criteria consisted of call rate > 0.95, MAF > 0.01, and Hardy-Weinberg equilibrium test statistic > 1E-4. Participants were excluded due to the following criteria: high-missingness (> 5% for genotyped variants), sex discordance, extreme heterozygosity (F statistics > 0.15), Non-European ancestry confirmed by joint analysis with HapMap 3 data using principal component (Outside of mean +/- 6 SD in PC1 or PC2 for European reference samples) (International HapMap 3 Consortium et al., 2010), and excessive relatedness (pairwise kinships > 0.125). We used PLINK version 1.9 for the above filtering (Purcell *et al.*, 2007).

For all samples and variants passing quality control, imputation was conducted for chromosome 1 to 22 using 1000 genome European reference panel at the Michigan Imputation Server (Das *et al.*, 2016) at the default setting, with the exception of the whole genome sequenced PPMI dataset. SNPs with an imputation quality of less than 0.3 and MAF < 1% were excluded. After quality control, the number of variants were approximately 2.6 - 2.9 million in NET-PD_LS1, DIGPD_neuroX, HBS, PDBP, and PRECEPT; 7.7 - 7.8 million in PICNICS, PROPARK, PARKWEST, DATATOP, PARKFIT, DIGPD, and OSLO; and 8.6 million in PPMI.

### Cohort-level analyses

We conducted a separate GWAS for each cohort per phenotype of interest. In addition, DIGPD cohorts were analyzed separately according to the genotyping array used (DIGPD_neuroX cohort and DIGPD_chip cohort). Each outcome was analyzed by an additive model with covariates. For the binomial outcomes at baseline, when the outcomes were positive for more than 5% of participants and >20 counts, logistic regression analyses were conducted. Those without the binomial outcome at baseline were followed-up until either censored or the development of the outcome. If more than 20 events were observed during follow-ups, the outcome was analyzed using cox proportional hazard models with time-varying covariates. For the analysis of continuous traits, linear mixed models were used to evaluate the variants’ association for the mean difference over time. Age at diagnosis, year from diagnosis to the observation, and sex were adjusted for in all analyses. In addition, the following covariates were associated with the outcome of interest in a backwards stepwise manner: quadratic age, quadratic years from diagnosis, years of education, medication status (levodopa usage, dopamine agonist usage, using either dopamine agonist or levodopa), and a Hoehn and Yahr score of 2 or more at the first observation (except for the models regressing for Hoehn and Yahr score itself or UPDRS motor score). These covariates were selected per study using Akaike’s Information Criteria (AIC) for logistic models and Cox survival models, and conditional AIC (cAIC) for linear mixed effect models. The cohort level analyses were conducted with R (version 3.5.0 https://www.r-project.org/) and rvtests (Zhan *et al.*, 2016). R package ‘cAIC4’ (Säfken *et al.*, 2018) was used to calculate cAIC.

### Meta-Analyses

The results from cohort-level analyses were combined using an inverse variance weighted fixed effect model. If the study-specific genomic inflation factor was more than 1.2, the study was excluded from the meta-analysis. Five of the 204 GWAS were excluded based on these criteria. For other cohorts, the overall alpha error was corrected using the genomic inflation factor before the meta-analysis. Meta-analyses were carried out with METAL (Willer *et al.*, 2010). From the meta-analysis results, we only evaluated variants with MAF > 0.05 due to statistical power constraints. We also excluded variants with minor allele frequency variability greater than 15% across cohorts. Further exclusions at the meta-analysis level include variants with Cochran’s Q-test for heterogeneity < 0.05 and a total participant N < 1000. The null hypothesis was tested with a significance level of 5E-8 on a two-sided test. For genome-wide signals, additional visualization and functional analyses were conducted using LocusZoom(Pruim *et al.*, 2010), FUMA (http://fuma.ctglab.nl/snp2gene/, version 1.3.3d) (Watanabe *et al.*, 2017). FUMA is a web-based annotation tool using MAGMA to conduct gene-based tests, a gene-set analysis and a tissue expression analysis. We applied a default setting. Also, we explored in eQTLGen database (http://www.eqtlgen.org/) (Võsa *et al.*, 2018) and meta-analyzed expression data in the brain accessible from the study by Qi et al.(Qi *et al.*, 2018).

Associations with the variants of interest, including the recently identified 90 risk variants for PD, known *LRRK2* and *GBA* variants, and *APOE*, were extracted from the meta-analysis results. We evaluated the associations of these variants and clinical features based on the significance level of 0.05, applying the Bonferroni adjustment of a maximum of 25 tests per variant (raw P-value < 0.002).

The summary of analytical processes is shown in figure 1.

**Figure 1:**
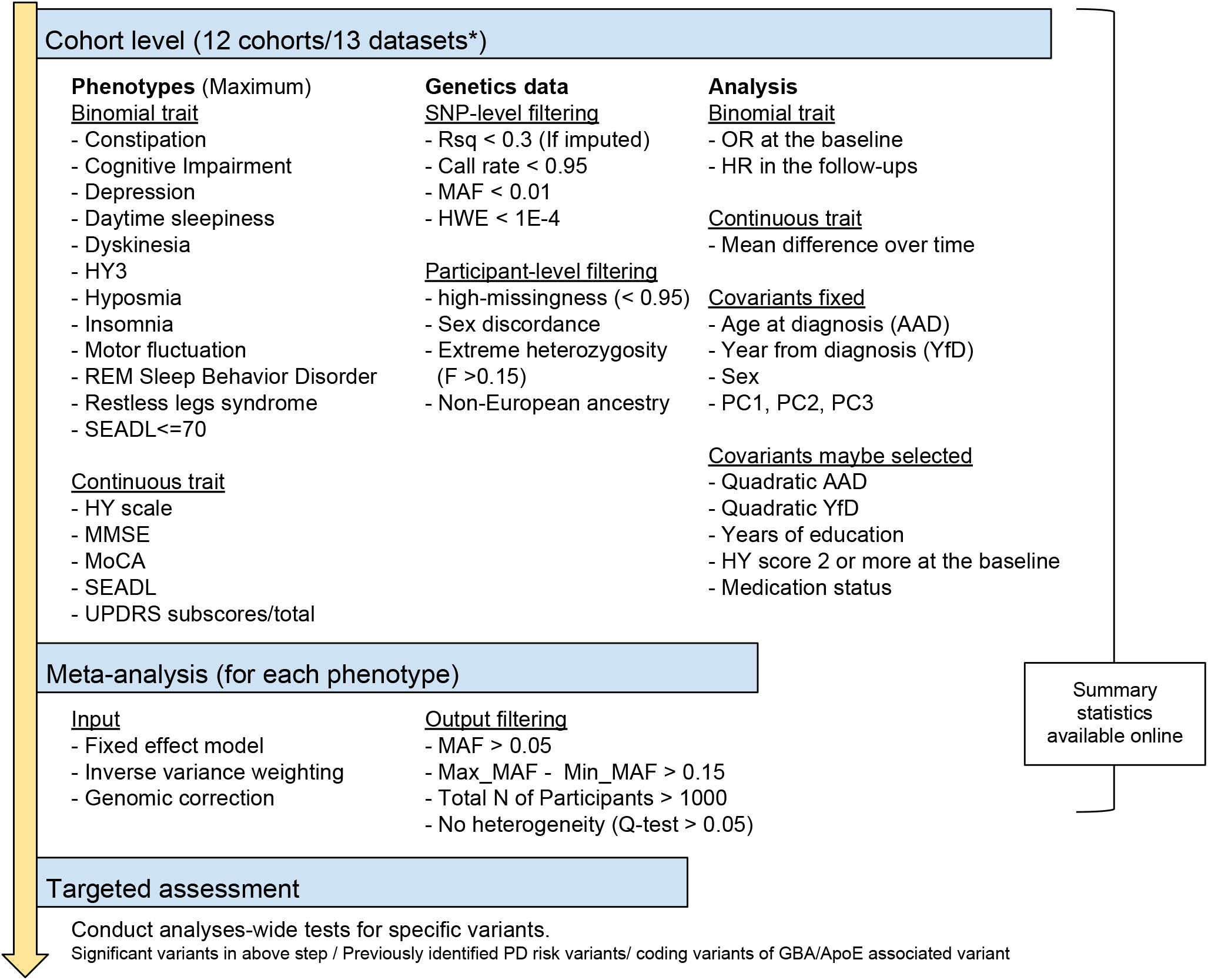
Graphical overview of the analysis strategy. * DIGPD cohort was analyzed separately depending on the genotyping system. Rsq, R square; MAF, Minor allele frequency; HWE, Hardy–Weinberg equilibrium test; OR, Odds ratio; HR, Hazard ratio; PC, Principal components; AAD, Age at diagnosis; YfD, Years from diagnosis to observation; HY score, the score on the Hoehn and Yahr scale;

## Results

### Novel GWAS associations with PD progression markers

The cohort characteristics are provided in Table 1. Overall, we analyzed 4,093 participants with 25,254 longitudinal data points over a median of 3.81 years. These cohorts varied in the years between enrollment and diagnosis, as well as follow-up durations. DATATOP, ParkWest, PPMI, and PreCEPT/PostCEPT enrolled untreated PD patients while others enrolled both treated and untreated patients. Considering the difference in design and recruitment strategies in the cohorts (Appendix), it is important to adjust for baseline characteristics as well as the follow-up lengths per cohort-level. All cohort-specific models for analysis are listed in Supplemental Table 2.

In total, 204 GWAS were conducted and combined into 33 meta-analyses. Eight meta-analyses were not evaluated because of the small number of total participants in the analyses (N total <1000). Those excluded were baseline analyses for RBD, RLS and SEADL70; and longitudinal analyses for constipation, daytime sleepiness, hyposmia, RBD, and RLS. Therefore, we investigated 9 binomial traits at baseline, 7 binomial traits for survival, and 9 continuous traits over the follow-ups. The genomic inflation factor was the mean value of 0.993, SD of 0.023, and the range was [0.951, 1.031] across meta-analyses. The summary statistics of the meta-analysis results, including the ones which were not evaluated in this manuscript, are publicly available for convenient browsing and downloading (https://pdgenetics.shinyapps.io/pdprogmetagwasbrowser/)

One association with the progression of PD was of genome-wide significance (P-value < 5.00E-08). The minor allele of rs382940 (chr9:108058562T>A), an intronic variant of *SLC44A1*, was associated with a higher hazard ratio (HR) of reaching Hoehn and Yahr stage 3.0 or greater (HR 2.04 [1.58, 2.62], P-value = 3.46E-8). When considering the baseline observations, the minor allele of rs61863020 (chr10:112956055G>A), an intergenic variant, was also significantly associated with the lower baseline OR of having insomnia (OR 0.63 [0,52, 0.75], P-value = 4.74E-8). Locus plots and forest plots for these two associations are shown in Figure 2. Cochran’s Q statistics, I-square and forest plots all showed no evidence of heterogeneity for these associations. (Figure 2)

**Figure 2:**
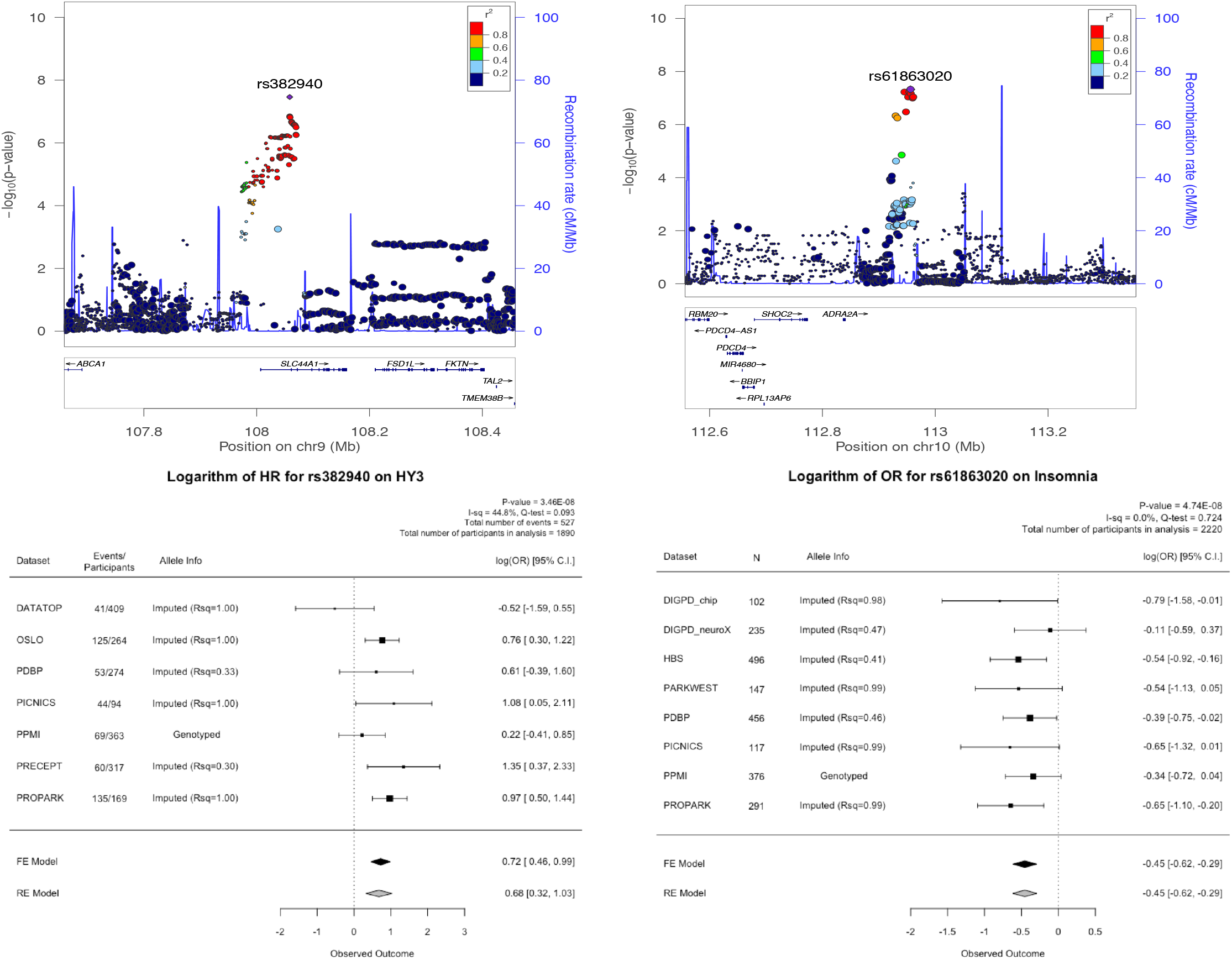
Locuszoom plots and forest plots of the two genome-wide significant hits. A: The locus plot for rs382940 which is associated with HY3. B: The locus plot for rs61863020 which is associated with insomnia. C: The forest plot for rs382940. D:The forest plot for rs61863020.

To evaluate the potential molecular mechanism for the two genome-wide signals, we explored eQTL datasets in blood and brain(Qi *et al.*, 2018; Võsa *et al.*, 2018), and functional annotation of the GWAS summary statistics using FUMA. Although it is in a regulatory region of *SLC44A*, rs382940 itself was not reported to be an eQTL in blood or brain. Gene-based tests using the GWAS summary statistics for reaching HY3 showed that *SLC44A1* was significant gene-wise(P-value = 5.8E-07 < Bonferroni correction threshold = 2.7E-6, supplemental figure 1). Rs61863020 was a significant eQTL for *ADRA2A* (α-2A adrenergic receptor) (P-value = 7.2E-4, the Bonferroni corrected P-value = 6.5E-3, up-regulation for A allele) in the brain.

In the meta-analysis results from the other clinical outcomes, rs382940 was associated with higher scores in the UPDRS part 2 and part 3 (UPDRS2_scaled: 0.36 [0.15, 0.57], P-value = 8.21E-04; UPDRS3_scaled: 0.29 [0.14, 0.45], P-value = 2.18E-04). These findings are consistent with the primary association of rs382940 and reaching HY3, which is a significant motor milestone (bilateral signs on clinical examination and the emergence of postural instability). Except for the association with having insomnia at baseline, rs61863020 was not significantly associated with other clinical variables in this analysis after adjusting for 25 tests.

### Targeted assessment for the PD risk variants

Of the 90 risk variants from the recently published PD GWAS, rs34637584 (*LRRK2* p.G2019S) and rs76763715 (GBA p.N370S) were not available in the meta-analyses because of their minor allele frequency (MAF) < 0.01. The remaining 88 PD GWAS risk SNPs were assessed in our 25 GWAS summary sets, resulting in evaluations of 2022 candidate associations. 112 associations between known genetic risk variants and clinical markers had raw p-values less than 0.05. After Bonferroni correction for all evaluated candidate associations, nine surpassed the threshold of the analyses-wide significance for the maximum of 25 analyses per variant (raw P-value < 0.002). The directions of these associations generally indicated that having the higher risk allele was associated with more severe deficits in both the cognitive and motor domains of PD, but not for sleeping problems. Having the risk allele (A) of rs1293298 (intron variant of *CTSB*) was associated with a lower risk of developing insomnia (HR 0.79 [0.69, 0.91], P-value = 1.2E-3), and the risk allele (A) of rs6500328 (intron variant of *NOD2*) and (A) of rs76116224 (intergenic variant close to 3’ end of *KCNS3)* were associated with a lower prevalence of daytime sleepiness at baseline (OR 0.76 [0.64, 0.90], P-value = 1.4E-3; OR 0.47 [0.32, 0.68], P-value = 8.4E-5; respectively). Among the 10 associations with analysis-wide significance, two were significant after adjusting for 88 variants (raw P-value < 5.68E-4), and one had test-wide significance (raw P-value < 2.47E-5). Figure 3 shows the strength of the associations for the selected variants with associations of analyses-wide significance in at least one analysis. This figure suggests that some risk variants were associated with specific clinical features. For example, rs35749011 was associated with both the HR of cognitive impairment at test-wide significance (HR 2.45 [1.64, 3.65] for the minor allele, P-value = 1.1E-5) and lower MoCA score over time at analyses-wide significance (−1.16 [−1.89, −0.43], P-value = 0.0018). Although it is an intergenic variant whose closest gene is *KRTCAP2*, the variant is in high LD (r2 = 0.78) with rs2230288 (*GBA* p.E365K) (Berge-Seidl *et al.*, 2017; Blauwendraat *et al.*, 2018), and has a similar spectrum of phenotype associations as rs2230288. Other notable variants with variant-wide significance were rs76904798, the intergenic variant close to the 5’ end of *LRRK2*, for reaching HY3 (HR 1.32 [1.14, 1.54] with the minor allele of T, P-value = 3.0E-4), and rs76116224 and the baseline OR of having daytime sleepiness mentioned above. The detailed information for all of the test results is provided as supplemental material. (Supplemental Table and Supplemental Figure)

**Figure 3:**
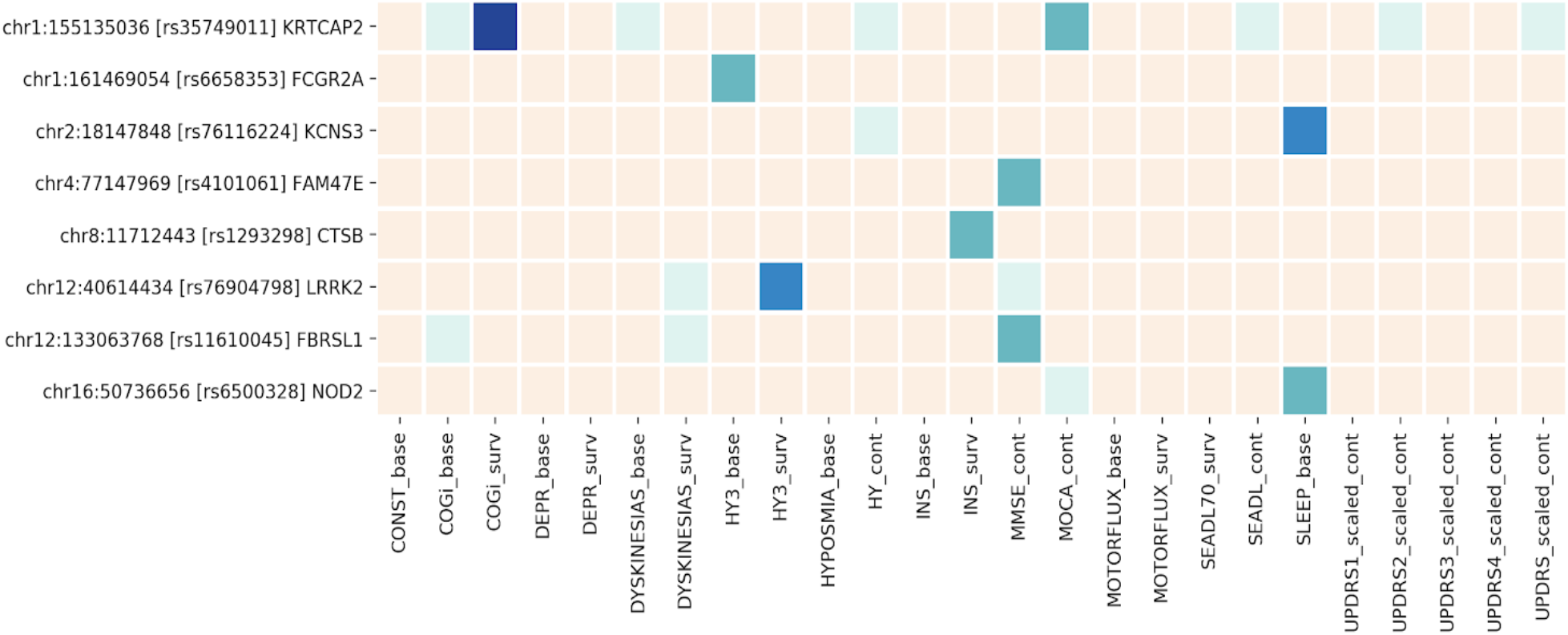
Heatmap of the Parkinson’s disease GWAS loci associated with progression markers. Cream, P-value > 0.05; light green, P-value < 0.05; green: P-value < 0.002; blue, P-value < 5.68E-4; dark blue, P-value < 2.47E-5). CONST, constipation; COGi, cognitive impairment; DEPR, depression; HY3, Hoehn and Yahr score; INS, insomnia; MMSE, Mini-Mental State Examination; MOCA, Montreal Cognitive Assessment; SEADL70, the modified Schwab and England Activities of Daily Living Scale; SLEEP, daytime sleepiness; UPDRS, Unified Parkinson’s Disease Rating Scale (UPDRS) or the Movement Disorder Society revised UPDRS, scaled at the baseline (UPDRS1-3) or during the course. Suffix of ‘base’ indicates the logistic regression model at baseline, ‘surv’ for the survival analysis over the course, and ‘cont’ for the mean difference overtime analyzed by linear mixed model.

**Figure 4:**
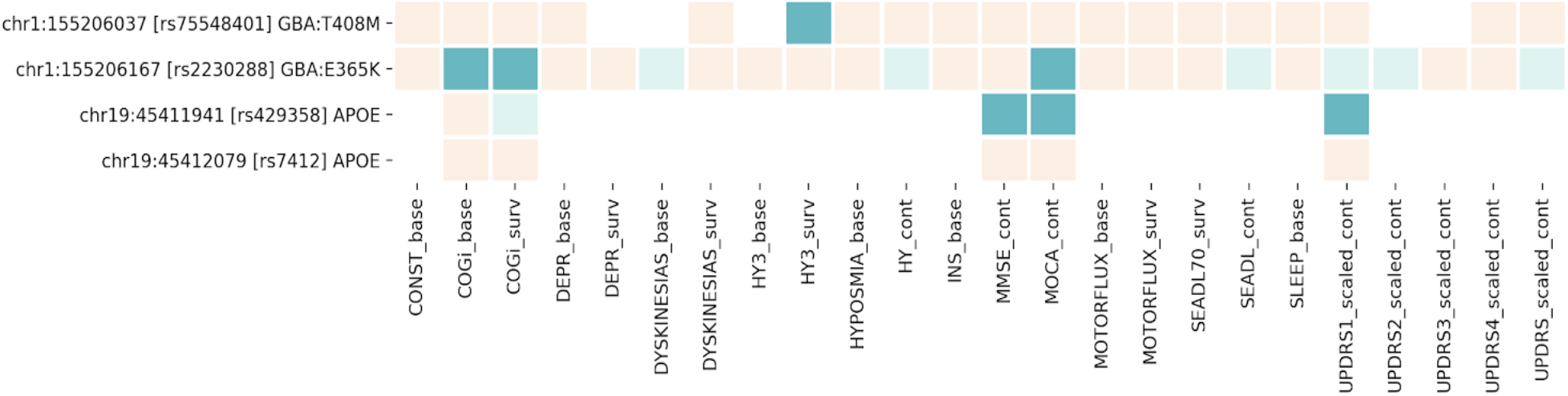
Heatmap of the GBA and APOE variants associated with progression markers. Cream, P-value > 0.05; light green, P-value < 0.05; green: Bonferroni corrected P-value < 0.05; CONST, constipation; COGi, cognitive impairment; DEPR, depression; HY3, Hoehn and Yahr score; INS, insomnia; MMSE, Mini-Mental State Examination; MOCA, Montreal Cognitive Assessment; SEADL, the modified Schwab and England Activities of Daily Living Scale; SLEEP, daytime sleepiness; UPDRS, Unified Parkinson’s Disease Rating Scale (UPDRS) or the Movement Disorder Society revised UPDRS, scaled at the baseline (UPDRS1-3) or during the course. Suffix of ‘base’indicates the logistic regression model at baseline, ‘surv’ for the survival analysis over the course, and ‘cont’ for the mean difference overtime analyzed by linear mixed model.

### *GBA* protein coding variants and APOE tagging variants

In the focused analyses for *GBA* coding variants, rs75548401, *GBA* p.T408M, was associated with the faster development of HY3 (HR 2.35 [1.58, 3.49], P-value = 2.5E-5). rs2230288, *GBA* p.E365K, was associated with the higher odds of having cognitive impairment at baseline (OR 2.05 [1.33, 3.18], P-value = 1.3E-3), faster development of cognitive impairment (HR 2.58 [1.71, 3.89] P-value = 5.5E-6), and lower MoCA score at the analysis-wide significance (Beta −1.23 [−1.97, −0.50], P-value = 1.0E-3). We previously reported these associations (under revision at *Neurology Genetics*) and we were able to confirm them in our updated analysis with more stringent multiple testing correction (FDR vs Bonferroni).

The C allele of rs429358, the tagging variant for the *APOE* E4 allele, was associated with lower MMSE (Beta-0.20 [−0.33, −0.07], P-value = 2.8E-3) and lower MoCA (Beta −0.52 [−0.86, −0.17], P-value = 3.4E-3) as expected. Moreover, it was associated with higher UPDRS part 1 scores (Beta 0.12 [0.04, 0.20] in Z score, P-value = 4.5E-3). We did not have enough evidence to conclude that the *APOE* E4 allele was associated with the prevalence of cognitive impairment at baseline (P-value = 0.4) or its development during follow-ups (P-value = 0.034). The T allele of rs7412 showed no association with these measurements, also predicted as this variant tagging *APOE* E2.

## Discussion

We conducted GWAS using longitudinal data from multiple PD cohorts to investigate markers of PD progression and heterogeneity. Of the 25 meta-analyses that we evaluated, we identified two variant-phenotype associations with genome-wide significance.

We also evaluated the summary statistics to assess clinical value of the variants of interest. One of our genome-wide hits, rs382940, in the intron of *SLC44A1*, was associated with a faster rate of progression to reach HY3. *SLC44A1*, soluble carrier 44A1, is also referred to as choline transporter-like protein 1 (CTL1). The gene is ubiquitously expressed in the brain, colon, thyroid and other organs and is involved in choline transport. No associations with PD and this variant or the gene itself have been reported so far although it has been studied in several vitro and vivo studies (Machová *et al.*, 2009; Schenkel *et al.*, 2015; Heffernan *et al.*, 2017; Gao *et al.*, 2018). Further investigation is warranted. The search of the Brain eQTL database suggested that another GWAS-signal, rs61863020, was associated with *ADRA2A* expression, a gene reported to be associated with arousal/sleep state (Gelegen *et al.*, 2014). *ADRA2A* is consistently expressed in locus coeruleus as well as nigral dopamine neurons and pyramidal neurons of the human brain (http://www.humanbraincode.org/, (Dong *et al*, 2018). The *ADRA2A*-encoded alpha2 adrenoreceptor modulates norepinephrine levels. Interestingly, norepinephrine (Tong *et al.*, 2006) and its receptors (Srinivasan and Schmidt, 2004; Mittal *et al.*, 2017) have been linked to PD in multiple model systems.

In the targeted assessments, we confirmed the previous results of the associations between *GBA* risk variants and motor and cognitive aspects of PD (Winder-Rhodes *et al.*, 2013; Brockmann *et al.*, 2015; Davis *et al.*, 2016a, b; Liu *et al.*, 2016). In contrast with *GBA* variants, association studies of *APOE* and cognitive function in PD have yielded mixed results (Huang *et al*, 2006; Kurz *et al.*, 2009; Federoff *et al.*, 2012; Mata *et al.*, 2014; Paul *et al.*, 2016). Our data supported the association of *APOE* and cognitive function on two measurements; MMSE and MoCA.

The strength of the current study is the hypothesis-free approach of GWAS, which can be powerful in identifying new associations and expanding our biological knowledge-base. While the associations here should be replicated and further investigated with vivo/vitro experiments, these findings suggest the prioritization of the two variants and loci for future validations. We have reported all of the summary results on our publicly accessible site to benefit researchers so that they may conduct/replicate the analysis of variants of interest in their own research.

The major limitation of this study, and studies like it, is the heterogeneity of the cohorts, which is apparent in several ways: baseline characteristics, definitions of binomial outcomes, patterns for clinical care over the course of follow-up, the platforms for genotyping/sequencing, and sample acquisition/enrollment practices. By meta-analyzing at the dataset-level and exercising careful quality control throughout, we tried to extract the most generalizable and reliable results across cohorts.

Another limitation is the power of the study. Although we have aggregated the largest collection of longitudinal data in PD genetics so far, more data would be needed to identify relatively small differences expected within PD patients compared to the case-vs-control setting.

Finally, the study participants were restricted to individuals with European ancestry. We are now striving to collect more data, including from populations that are under-represented in this study, to improve our understanding of this topic in future studies.

## Conclusion

With 4,093 participants and 25,254 longitudinal data points over a median of 3.81 years, we performed 25 GWAS meta-analyses. We found two genome-wide significant signals: the rate to reach HY3 during the disease course and rs382940; and the prevalence of insomnia at baseline and rs61863020. We also conducted targeted assessments of previously published variants of interest using the GWAS results. These results provide valuable insights into how genetic factors contribute to the heterogeneity of PD and disease progression.

## Supplementary Materials

Appendix: The description of study cohorts

Supplemental table 1: Cohort specific definitions of binomial outcomes

Supplemental table 2: Analytical models per datasets

Supplemental table 3: The meta-analysis results of the association between risk variants and the clinical features and progression of Parkinson’s disease

Supplemental figure 1: Gene-based test for reaching Hoehn and Yahr stage 3 or higher

Supplemental figure 2: Heatmap for the meta-analysis results of the association between risk variants and the clinical features and progression of Parkinson’s disease

