## Appendix for "Genome-wide association study of Parkinson’s disease progression biomarkers in 12 longitudinal patients’ cohorts"

**Appendix:** **The description of study cohorts**

Deprenyl and Tocopherol Antioxidative Therapy of Parkinsonism (DATATOP**)** was a randomized clinical trial conducted between September 1987 and November 1989 at 28 sites across US and Canada. The primary objective was to test the efficacy of deprenyl and/or tocopherol. 800 patients with Parkinson’s disease diagnosed within 5 years and not requiring symptomatic treatment were observed for up to 2 years.^1^ The study was supported by a Public Health Service grant (NS24778) from the National Institute of Neurological Disorders and Stroke; by grants from the General Clinical Research Centers Program of the National Institutes of Health at Columbia University (RR00645), the University of Virginia (RR00847), the University of Pennsylvania (RR00040), the University of Iowa (RR00059), Ohio State University (RR00034), Massachusetts General Hospital (RR01066), the University of Rochester (RR00044), Brown University (RR02038), Oregon Health Sciences University (RR00334), Baylor College of Medicine (RR00350), the University of California, San Diego (RR00827), Johns Hopkins University (RR00035), the University of Michigan (RR00042), and Washington University (RR00036); the Parkinson's Disease Foundation at Columbia-Presbyterian Medical Center, New York; the National Parkinson Foundation, Miami; the Parkinson Foundation of Canada, Toronto; the United Parkinson Foundation, Chicago; the American Parkinson's Disease Association, New York; and the University of Rochester, Rochester, N.Y.

Drug Interaction with Genes in Parkinson's Disease (DIGPD**)** is a cohort with 413 patients with Parkinson’s disease diagnosed by UK Parkinson’s disease society brain bank clinical diagnostic (UKPDSBB) criteria with disease duration less than 5 years at the entry.^2^ It is an ongoing study since 2009, and the participants are followed for up to 7 years at eight sites in France. (Corvol et al., in press in Neurology). DNA samples were collected from all of them. DIGPD is sponsored by Assistance Publique Hôpitaux de Paris, funded by a grant from the French Ministry of Health (PHRC 2008, AOM08010) and a grant from the Agence Nationale pour la Sécurité des Médicaments (ANSM 2013).

Harvard Biomarkers Study (HBS) is a longitudinal case-control study. More than 2,700 individuals with early-stage PD, patients with memory impairment, and controls without neurological disease were enrolled and longitudinally phenotyped since 2008.^3^ HBS was supported by the Harvard NeuroDiscovery Center, MJFF, NINDS U01NS082157, U01NS100603, and the Massachusetts Alzheimer’s Disease Research Center NIA P50AG005134.

NIH Exploratory Trials in Parkinson's Disease Large Simple Study 1 (NET-PD LS1**)** was a randomized study conducted between March 2007 and September 2013 to determine if the nutritional supplement creatine slows the clinical progression of Parkinson’s disease over time. 1741 patients from 50 sites in the US and Canada participated.^4^ They were within 5 years from diagnosis. The plan was for them to be followed for at least 5 years, but the study ended early for futility based on an interim analysis at which point the median follow-up time was 4 years. Financial support for the LS-1 study was provided by National Institute of Neurological Disorders and Stroke (NINDS) grant U01NS43128.

Oslo PD study[[Citation error]](http://127.0.0.1:8080/c/error) (Oslo**)** is an ongoing study since 2007, with 317 patients diagnosed with ULPDSBB criteria with modification of allowing family history. The participants are being followed up to 6 years in prospective (30 years in retrospective) at Oslo University Hospital in Norway.^5^ Oslo PD is supported by the Research Council of Norway and South-Eastern Norway Regional Health Authority.

ParkFit cohort was originally a randomized trial evaluating a multifaceted behavioural change programme to increase physical activities in patients with Parkinson’s disease.^6^ The study conducted from September 2008 to February 2012 at a single center in the Netherlands, with 586 patients with Parkinson’s disease diagnosed by UKPDSBB, with Hoehn Yahr stage 3 or lower, and with sedentary lifestyle at the entry. They were followed up for 2 years. The primary objective was concluded as not significant^6^. ParkFit is supported by ZonMw (the Netherlands Organization for Health Research and Development (75020012)) and the Michael J Fox Foundation for Parkinson’s research, VGZ (health insurance company), GlaxoSmithKline, and the National Parkinson Foundation.

The Norwegian ParkWest study (ParkWest) is an ongoing prospective longitudinal multicenter cohort study of patients with incident Parkinson’s disease from Western and Southern Norway, designed to study the incidence, neurobiology and prognosis of PD.^7^ Between November 1st 2004 and 31st of August 2006, all new cases of Parkinson Disease within the study area (Sogn and Fjordane, Hordaland, Rogaland and Aust-Agder) were recruited, and since the start of the study 212 of these patients and their age-/sex-matched control group were followed. The Norwegian ParkWest study is supported by the Research Council of Norway, the Western Norway Regional Health Authority, Stavanger University Hospital Research Funds, and the Norwegian Parkinson’s Disease Association.

The National Institute of Neurological Disorders and Stroke (NINDS) Parkinson’s Disease Biomarker Program (PDBP) is aiming to discover new diagnostic and progression biomarkers for Parkinson’s disease.^8^ It is a combined cohort of 9 PDBP-funded research studies. The members have various stages of Parkinson’s disease and recruited throughout the United States.

Parkinsonism: Incidence and Cognitive and Non-motor heterogeneity In Cambridgeshire (PICNICS) is a population-based longitudinal study of 282 incident PD cases recruited between 2008 and 2013 with ongoing follow-up at 18 month intervals.^9,10^ PD cases were diagnosed based on the UKPDSBB criteria, and followed up at a single center in the UK. PICNICS has received funding from the Cure Parkinson’s Trust, the Van Geest Foundation and is supported by the the National Institute of Health Research Cambridge Biomedical Research Centre.

Parkinson’s progression markers initiative (PPMI) is an ongoing study started on July 2010, enrolling 424 patients with Parkinson’s disease diagnosed within 2 years from the study entry date.^11^ The study sites are located in 33 sites across the US, Europe, Israel and Australia^11^. PPMI is supported by the Michael J Fox Foundation for Parkinson’s Research.

Parkinson Research Examination of CEP1348 Trial **(**PreCEPT**)** is a clinical trial of the mixed lineage kinase inhibitor CEP‐1357,4 sponsored by Cephalon, Inc. (West Chester, PA) and H. Lundbeck A/S (Valby-Copenhagen, Denmark). The study was conducted at 65 sites in North America. The trial enrolled 806 early, untreated PD patients within one year from the onset. The original trial was started in April 2002 and terminated in August 2005 due to the futility, but the participants were continuously followed-up in the prospective observational study (PostCEPT).^12^

The studies were funded by NINDS 5U01NS050095‐05, Department of Defense Neurotoxin Exposure Treatment Parkinson's Research Program. Grant Number: W23RRYX7022N606, the Michael J Fox Foundation for Parkinson’s research, Parkinson's Disease Foundation, Lundbeck Pharmaceuticals. Cephalon Inc, Lundbeck Inc, John Blume Foundation, Smart Family Foundation, RJG Foundation, Kinetics Foundation, National Parkinson Foundation, Amarin Neuroscience LTD, CHDI Foundation Inc, National Institutes of Health (NHGRI, NINDS), Columbia Parkinson's Disease Research Center.

Profiling Parkinson’s disease study (ProPark) is an ongoing study started from May 2003. Initially, 420 patients recruited in several sites in the Netherlands by March 2006.^13^ Patients were diagnosed with UKPDSBB criteria and in various disease durations at the enrollment. They are evaluated annually with the SCOPA scale. This study is funded by the Alkemade-Keuls Foundation, Stichting Parkinson Fonds, Parkinson Vereniging, The Netherlands Organisation for Health Research and Development.

### **Study investigators**

Harvard Biomarkers Study. Co-Directors: Harvard NeuroDiscovery Center: Clemens R. Scherzer, Bradley T. Hyman, Charles G. Jennings; Investigators and Study Coordinators: Harvard NeuroDiscovery Center: Yuliya Kuras, Daly Franco, Frank Zhu; Brigham and Women’s Hospital: Lewis R. Sudarsky, Michael T. Hayes, Chizoba C. Umeh, Reisa Sperling; Massachusetts General Hospital: John H. Growdon, Michael A. Schwarzschild, Albert Y. Hung, Alice W. Flaherty, Deborah Blacker, Anne-Marie Wills, U. Shivraj Sohur, Vivek K. Unni, Nicte I. Mejia, Anand Viswanathan, Stephen N. Gomperts, Vikram Khurana, Mark W. Albers, Maria Allora-Palli, Alireza Atri, David Hsu, Alexandra Kimball, Scott McGinnis, Nutan Sharma, John Becker, Randy Buckner, Thomas Byrne, Maura Copeland, Bradford Dickerson, Matthew Frosch, Theresa Gomez-Isla, Steven Greenberg, James Gusella, Julius Hedden, Elizabeth Hedley-Whyte, Keith Johnson, Raymond Kelleher, Aaron Koenig, Maria Marquis-Sayagues, Gad Marshall, Sergi Martinez-Ramirez, Donald McLaren, Olivia Okereke, Elena Ratti, Christopher William, Koene Van Dij, Shuko Takeda, Anat Stemmer-Rachaminov, Jessica Kloppenburg, Catherine Munro, Rachel Schmid, Sarah Wigman, Sara Wlodarcsyk; University of Ottawa: Michael G. Schlossmacher; Scientific Advisory Board: Massachusetts General Hospital: John H. Growdon; Brigham and Women’s Hospital: Dennis J. Selkoe, Reisa Sperling; Harvard School of Public Health: Alberto Ascherio; Data Coordination: Harvard NeuroDiscovery Center: Thomas Yi, Massachusetts General Hospital: Joseph J. Locascio, Haining Li; Biobank Management Staff: Harvard NeuroDiscovery Center: Gabriel Stalberg, Zhixiang Liao.

Parkinson Study Group DATATOP investigators: *Steering Committee* — Ira Shoulson, M.D. (principal investigator), University of Rochester, Rochester, N.Y.; Stanley Fahn, M.D. (co-principal investigator), Columbia–Presbyterian Medical Center, New York; David Oakes, Ph.D. (chief biostatistician, 1987 to present), University of Rochester; Charles Odoroff Ph.D. (deceased) (chief biostatistician, 1985–1987), University of Rochester; Anthony Lang, M.D., Toronto Western Hospital, Toronto; J. William Langston, M.D., California Parkinson's Foundation, San Jose, Calif.; Peter LeWitt, M.D., Sinai Hospital, Detroit; Warren Olanow, M.D., University of South Florida, Tampa; John B. Penney, M.D. (deceased), University of Michigan, Ann Arbor; and Caroline Tanner, M.D., Rush–Presbyterian–St. Luke's Medical Center, Chicago. *Participating Investigators* — William Koller, M.D. (deceased), University of Kansas, Kansas City; Warren Olanow, M.D., University of South Florida; Robert Rodnitzky, M.D., University of Iowa, Iowa City; J. Stephen Fink, M.D., Ph.D. (deceased), and John H. Growdon, M.D., Massachusetts General Hospital, Boston; George Paulson, M.D., Ohio State University, Columbus; Roger Kurlan, M.D., University of Rochester; Joseph H. Friedman, M.D., Roger Williams General Hospital, Providence; Stephen Gancher, M.D., and John Nutt, M.D., Oregon Health Sciences University, Portland; Ali H. Rajput, M.D., University of Saskatchewan, Saskatoon; James Bennett, M.D., Ph.D., and George F. Wooten, M.D., University of Virginia, Charlottesville; Peter LeWitt, M.D., Sinai Hospital, Detroit; Christopher Goetz, M.D., Caroline Tanner, M.D., Kathleen Shannon, M.D., and Harold Klawans, M.D. (deceased), Rush–Presbyterian–St. Luke's Medical Center, Chicago; Oksana Suchowersky, M.D., University of Calgary, Calgary, Alta.; Mitchell Brin, M.D., and Susan Bressman, M.D., Columbia–Presbyterian Medical Center, New York; William J. Weiner, M.D. (deceased), and Juan Sanchez-Ramos, M.D., Ph.D., University of Miami, Miami; Joseph Jankovic, M.D., Baylor College of Medicine, Houston; John B. Penney, M.D. (deceased), University of Michigan, Ann Arbor; Anthony Lang, M.D., Toronto Western Hospital, Toronto; Margaret Hoehn, M.D. (deceased), St. Luke's Hospital, Denver; James Tetrud, M.D., California Parkinson's Foundation, San Jose; J. David Grimes, M.D. (deceased), Ottawa Civic Hospital, Ottawa, Ont.; Ronald Pfeiffer, M.D., University of Nebraska and Creighton University, Omaha; Cliff Shults, M.D. (deceased), and Leon Thal, M.D. (deceased), University of California, San Diego; Serge Gauthier, M.D., Montreal General Hospital and McGill University, Montreal; Lawrence I. Golbe, M.D., University of Medicine and Dentistry of New Jersey, New Brunswick; Joel S. Perlmutter, M.D., Washington University, St. Louis; Hamilton Moses III, M.D., Johns Hopkins University, Baltimore; and Howard I. Hurtig, M.D., and Matthew Stern, M.D., The Graduate Hospital, Philadelphia.

*Site Coordinators* — Ruth Barter, R.N., and Bridget Vetere-Over-field, R.N., Kansas City, Kans.; Lisa Gauger, B.A., and Terresita Malapira, R.N., Tampa, Fla.; Judith Dobson, R.N., Iowa City, Iowa; Susan Atamian, R.N., Marsh Tennis, R.N., Jennifer B. Cohen, B.A., and Gena Desclos, B.A., Boston; Lena Denio, M.T., Steven Huber, Ph.D., and Teresa Woike, R.N., Columbus, Ohio; Jill Behr, R.N., M.S., and Irenita Gardiner, R.N., Rochester, N.Y.; Margaret Lannon, R.N., M.S., Providence, R.I.; Julie Carter, R.N., and Susanne Northrup, Portland, Ore.; Bernice Kanigan, R.N., Saskatoon, Sask.; Margaret Turk, R.N., M.S., and Elke Landow, R.N., Charlottesville, Va.; Patricia Schlick, R.N., and Kathie Mistura, R.N., Detroit; V. Susan Carrol, R.N., M.S., Jeana Thelen, R.N., and Joan Lechner, Chicago; Carol Demong, R.N., Calgary, Alta.; Linda Winfield, R.N., and Carol Moskowitz, R.N., New York; Angela Ingenito, R.N., Carol Sheldon, R.N., and Lisa Cornelius, B.A., Miami; Dorothy Heiberg, R.N., Houston; Jan Brady, R.N., M.S., Ann Arbor, Mich.; Catherine Kierans, R.N., M.A., and Loretta Bell-Scantlebury, R.N., Toronto; Helena Weber, M.T., M.A., Denver; Deborah Savoini, R.N., Paula Lewis, R.N., and S. Jerome Kutner, Ph.D., San Jose, Calif.; Peggy Gray, R.N., Ottawa, Ont.; Ruth Hofman, R.N., and Carolyn Glaeske, R.N., Omaha; Mary Margaret Pay, R.N., and David Salmon, Ph.D., San Diego; Frances McFaul, R.N., and Donna Amyot, R.N., Montreal; Mary Bergen, R.N., New Brunswick, N.J.; Lori McGee-Minnich, R.N., St. Louis; Patricia O'Donnell, R.N., M.S., Baltimore; and Susie Ferrise, R.N., and Kathy Shallow, B.A., Philadelphia.

*Coordination and Data Center* (University of Rochester Medical Center, Suite 160, 1325 Mt. Hope Ave., Rochester, NY 14620) — Rita M. Pelusio, M.S.Ed. (program manager); Alice Rudolph, Ph.D. (chief study coordinator); Peter Como, Ph.D. (neuropsychology consultant); Charlyne Miller, R.N., M.S. (nurse clinician); Michael Linsner, B.S., Joseph Connorton, B.A., and Judith Nusbaum, B.A. (analyst programmers); Carrie Irvine, B.S., and Constance Orme, B.A. (information analysts); Ruth Nobel, Deborah Baker, Donna LaDonna, Mary Ellen Rothfuss, and Lynn Doerr (deceased) (secretarial staff); Jacqueline Wendel, B.T. (CLINFO manager); and Belinda Rodriguez, Virginia Collins, Scott Dalston, and Paul Bivrell (student clerks).

*Biostatistics Center* (Division of Biostatistics, University of Rochester Medical Center, Rochester, NY 14642) — Charles Odoroff, Ph.D. (deceased), and David Oakes, Ph.D. (chief biostatisticians); Michael McDermott, Ph.D., and Shirley Eberly, M.S. (biostatisticians); Sandra Plumb, B.S. (lead programmer); Arthur Watts, B.S., Lori Yorkey, B.A., Anna Choi, B.A., and Karen Gerwitz, B.S. (analyst programmers).

*Pharmacy Center* (Strong Memorial Hospital, Rochester, NY 14642) — Paul Evans, R.Ph. (chief pharmacist); Lori Dellapena and Verna Singletary (pharmacy technicians).

*Safety Monitoring Committee* (Rochester, N.Y.) — Robert Herndon, M.D. (chair, January 1, 1987–June 30, 1988), Pierre Tariot, M.D. (chair, July 1, 1988 to present), Edward Bell, Pharm.D., Robert C. Griggs, M.D., W. Jackson Hall, Ph.D., Sandra Plumb, B.S. (lead programmer), and Arthur Watts, B.S. (analyst programmer).

*Scientific Advisory Committee* — C. David Marsden, D.Sc. (deceased) (chair), London; Gerald Cohen, Ph.D. (deceased), Joseph Fleiss, Ph.D. (deceased), and Richard Mayeux, M.D., New York; and Laurence Jacobs, M.D., and Arthur J. Moss, M.D., Rochester, N.Y.

*Clinical Trials Monitoring Committee* (National Institute of Neurological Diseases and Stroke) — Emanuel M. Stadlan, M.D. (chair), Bethesda, Md.; Milton Alter, M.D. (deceased), Philadelphia; Jesse Cedarbaum, M.D., White Plains, N.Y.; Jonas Ellenberg, Ph.D., Bethesda, Md.; and Robert Kibler, M.D., Atlanta.

*Assay Standards Committee* — Robert Roth, Ph.D. (chair), New Haven, Conn.; Harvey Cohen, M.D., Ph.D., Rochester, N.Y.; Matthew Galloway, Ph.D., Detroit; Ian Irwin, Ph.D., San Jose, Calif.; Peter LeWitt, M.D., Detroit; Govind Vatassery, Ph.D., Minneapolis.

*Neuropsychological Testing Committee* — Richard Mayeux, M.D. (chair), New York; Peter Como, Ph.D., Rochester, N.Y.; Jean St. Cyr, Ph.D., Toronto; Yaakov Stern, Ph.D., and Janet Williams, D.S.W., New York; and Robert Wilson, Ph.D., Chicago.

*Cerebrospinal Fluid Assay Center* (Wayne State University, Detroit, MI 48207) — Matthew P. Galloway, Ph.D. (director); Mike Kaplan (deceased) and Rashid Lodhi.

*Deprenyl Metabolities Assay Center* (Institute for Medical Research, San Jose, CA 95128) — Ian Irwin, Ph.D. (director).

*Blood Tocopherol Assay Center* (Our Lady of Mercy Medical Center, Bronx, NY 10466) — Edward Norkus, Ph.D. (director).

*Specimen Repository* (Department of Neurology, University of Rochester, Rochester, NY 14642) — Dorothy Flood, Ph.D. (director), Thomas McNeill, Ph.D., Norma Harary, Ph.D., and Laurie Koek, B.S.

*Laboratory Surveillance Testing* (SciCor Laboratories, Indianapolis, IN 46241) — Robert L. Creveling, M.D. (director).

Parkinson Study Group PRECEPT investigators: Steering committee: *University of Rochester, Rochester, NY*: Ira Shoulson, MD (principal investigator), Steven Schwid, MD (medical monitor), Christopher Hyson, MD (medical monitor), David Oakes, PhD (chief biostatistician), Emily Gorbold, BA, (project coordinator), Alice Rudolph, PhD (project coordinator), Aileen Shinaman, JD (Parkinson Study Group executive director), Cornelia Kamp, MBA (director, Clinical Research Operations), Karl Kieburtz, MD, MPH (director, Clinical Trials Coordination Center); *Toronto Western Hospital, University Health Network, Toronto, Ontario, Canada*: Anthony Lang, MD (coprincipal investigator); *Columbia University Medical Center, New York, NY*: Stanley Fahn, MD; *Duke University Medical Center, Durham, NC*: Lisa Gauger, BA; *Rush University Medical Center, Chicago, IL*: Christopher Goetz, MD; *Institute for Neurodegenerative Disorders, New Haven, CT*: Kenneth Marek, MD, John Seibyl, MD.

Participating investigators and coordinators: *Colorado Neurological Institute, Englewood, CO*: Lauren Seeberger, MD, Rajeev Kumar, MD; *London Health Sciences Center, London, Canada*: Mandar Jog, MD, Cheryl Horn, RN; *Rush–Presbyterian–St. Luke’s Medical Center, Chicago, IL*: Kathleen Shannon, MD, Lucia Blasucci, RN, CCRC; *University of Colorado Health Sciences Center, Denver, CO*: Maureen Leehey, MD, Teresa Derian, RN; *Ottawa Hospital Civic Site, Ottawa, Ontario, Canada*: David Grimes, MD, Melodie Mortensen, BSCN, Keely Haas, RN; *University of Minnesota/ Minnesota VA Medical Center, Minneapolis, MN*: Paul Tuite, MD, Susan Rolandelli, RN; *University of California Irvine, Irvine, CA*: Neal Hermanowicz, MD, Shari Niswonger, RN; *University of Rochester, Rochester, NY*: Roger Kurlan, MD, Irenita Gardiner, RN, CCRC; *Toronto Western Hospital, University Health Network, Toronto, Ontario, Canada*: Janis Miyasaki, MD, FRCPC, Lisa Johnston, RN, BSCN, CNN; *The Parkinson’s Institute, Sunnyvale, CA*: James Tetrud, MD, Tracy Stewart, RN; *NeuroHealth PD Movement Disorders Center, Warwick, RI*: Joseph Friedman, MD, Hubert Fernandez, MD, Margaret Lannon, RN, MS; *University of Iowa, Iowa City, IA*: Robert Rodnitzky, MD, Judith Dobson, RN, CCRC; *Mayo Clinic Arizona, Scottsdale, AZ*: Virgilio Evidente, MD, Marlene Lind, RN; *Oregon Health & Science University, Portland, OR*: Julie Carter, RN, MN, ANP, Pamela Andrews, BS; *Chum-Hotel Dieu/McGill Center for Studies in Aging, Montreal, Quebec, Canada*: Michel Panisset, MD; *Washington University School of Medicine, St. Louis, MO*: Brad Racette, MD, Patricia Deppen, RN; *Baylor College of Medicine, Houston, TX*: Joseph Jankovic, MD, Christine Hunter, RN, CCRC; *Institute for Neurodegenerative Disorders, New Haven, CT*: Danna Jennings, MD, Barbara Fussell, RN; *Albany Medical College, Albany, NY*: Eric Molho, MD, Stewart Factor, MD; *Indiana University School of Medicine, Indianapolis, IN*: Joanne Wojcieszek, MD; *University of California–Davis, Sacramento, CA*: Lin Zhang, MD, PhD, Lisa Wilson, MS, CCRP, Teresa Tempkin, RNC, MSN; *Duke University Medical Center, Durham, NC*: Burton Scott, MD, Joanne Field, BSN, RN; *Cleveland Clinic, Cleveland, OH*: Thyagarajan Subramanian, MD, Ruth Kolb, CCRP; *University of Pennsylvania, Philadelphia, PA*: Andrew Siderowf, MD, Amy Colcher, MD, Heather Maccarone, RN, BSN; *University of South Florida, Tampa, FL*: Robert Hauser, MD, Joanne Nemeth, RN; *Johns Hopkins University, Baltimore, MD*: Joseph Savitt, MD, PhD, Kevin Biglan, MD, MPH, Melissa Gerstenhaber, RNC, MSN; *University of Cincinnati/Cincinnati Children’s Hospital, Cincinnati, OH*: Alok Sahay, MD, Arif Dalvi, MD, Maureen Gartner, RN, Donna Schwieterman, MA, CCRC; *Mayo Clinic Jacksonville, Jacksonville, FL*: Ryan Uitti, MD, Margaret Turk, RN; *University of Sherbrooke, Sherbrooke, Quebec, Canada*: Jean Rivest, MD, Daniel Soucy, RN; *University of Virginia, Charlottesville, VA*: Frederick Wooten, MD, Elke Rost-Ruffner, RN, BSN; *Massachusetts General Hospital, Boston, MA*: Michael Schwarzschild, MD, PhD, Marsha Tennis, RN; *Medical College of Georgia, Augusta, GA*: Kapil Sethi, MD, Lisa Hatch, RN, BSN; *University of Tennessee–Memphis, Memphis, TN*: Ronald Pfeiffer, MD, Brenda Pfeiffer, RN, BSN; *North Shore–LIJ Health System, Manhasset, NY*: Andrew Feigin, MD, Jean Ayan, RN, Barbara Shannon, RN; *Northwestern University, Chicago, IL*: Tanya Simuni, MD, Karen Williams, BA, Michele Wolff, BA; *Medical University of Ohio, Toledo, OH*: Lawrence Elmer, MD, PhD, Kathy Davis, RN; *University of Connecticut, Glastonbury, CT*: Antonelle de Marcaida, MD, Sheila Thurlow, RN; *Hotel-Dieu Hospital–CHUM, Montreal, Quebec, Canada*: Sylvain Chouinard, MD, Hubert Poiffaut, RN; *Barrow Neurological Institute, Phoenix, AZ*: Holly Shill, MD, Mark Stacy, MD, Lynn Marlor, BSN, MSHS, Jill Danielson, RN; *The Parkinson’s & Movement Disorder Institute, Fountain Valley, CA*: Daniel Truong, MD, Jacky Vo, MS; *LSU Health Science Center Shreveport, Shreveport, LA*: Richard Zweig, MD, Rhonda Feldt, RN; *Columbia University Medical Center, NY, NY*: Cheryl Waters, MD, Angel Figueroa, BBA, Anne Tam, BS, CCRC; *University of Kansas Medical Center, Kansas City, KS*: Rajesh Pahwa, MD, Amy Parsons, RN, BSN; *University of Southern California, Los Angeles, CA*: Jennifer Hui, MD, Allan Wu, MD, Connie Kawai, RN, BSN, CCRC; *University of Alberta, Edmonton, AB, Canada*: Richard Camicioli, MD, Pamela King, BScN, RN; *University of Chicago, Chicago, IL*: Arif Dalvi, MD, Un Jung Kang, MD, Elizabeth Shaviers, Barbara Harding-Clay, CMA, CCRC; *University of Maryland School of Medicine, Baltimore, MD*: Stephen Reich, MD, Lisa Shulman, MD, Carol Dignon, RN, MSN, Kelly Dustin, RN: *UMDNJ Robert Wood Johnson Medical School, New Brunswick, NJ*: Margery Mark, MD, Deborah Caputo, RN, MSN; *Saskatoon Dist Health Board Royal University Hospital, Saskatoon SK, Canada*: Ali Rajput, MD; *Boston University, Boston, MA*: Peter Novak, MD, Cathi-Ann Thomas, RN, MS; *Pacific Neuroscience Medical Group, Oxnard, CA*: James Sutton, MD, Juanita Young, CCRC; *University of California–San Diego, La Jolla, CA*: David Song, MD, Deborah Fontaine, RNCS, MS; *Creighton University, Omaha, NE*: John M. Bertoni, MD, PhD, Carolyn Peterson, RN; *Medical College of Wisconsin, Milwaukee, WI*: Karen Blindauer, MD, Jeannine Petit, ANP; *Scott & White Hospital/Texas A&M University, Temple, TX*: Bala Manyam, MD, Danielle McNeil-Keller, LMSW, Jacqueline Whetteckey, MD; *Clinical Neuroscience Center, Southfield, MI*: Peter LeWitt, MD, Maryan DeAngelis, RN, CCRC; *University of Calgary, Calgary, AB, Canada*: Ranjit Ranawaya, MD, Oksana Suchowersky, MD, Carol Pantella, RN; *Brigham&Women’s Hospital, Boston, MA*: Lewis Sudarsky, MD; *Beth Israel Deaconess Medical Center, Boston, MA*: Daniel Tarsy, MD, Linda Paul, NP, Lisa Scollins, NP; *Long Island Jewish Medical Center, New Hyde Park, NY*: Mark Forrest Gordon, MD; *Beth Israel Medical Center, NY,* *NY*: Susan Bressman, MD, Alessandro DiRocco, MD, Karyn Boyar, RN, CNS, FNP; *Stanford University Medical Center, Stanford, CA*: Helen Bronte-Stewart, MD, Amy Andrzejewski, BS; *UMDNJ School of Osteopathic Medicine, Stratford, NJ*: Gerald Podskalny, DO; *Cleveland Clinic Florida–Weston, Weston, FL*: Nestor Galvez- Jimenez, MD, Jose Alvarez, CCRC; *University of Arkansas for Medical Sciences, Little Rock, AR*: Sami Harik, MD, Samer Tabbal, MD, Jana Patterson, RN

Biostatistics and coordination center staff: *University of Rochester, Rochester, NY*: Arthur Watts, BS, Rory Doolan, BA, Michele Goldstein, BS, Connie Orme, BA, Larry Preston, BPS, Tina Winebrenner.

Data monitoring committee: *The Parkinson’s Institute, Sunnyvale, CA*: Caroline Tanner, MD, PhD (chair); *Johns Hopkins University, Baltimore, MD*: Steven Piantadosi, MD, PhD; *University of British Columbia, Vancouver, BC, Canada*: Jon Stoessl, MD, Paul Keown, MD; *University of Pennsylvania, Philadelphia, PA*: Lynn Schuchter, MD.

The following employees of Cephalon, Inc. and H. Lundbeck A/S were substantively involved in the design, conduct, and analysis of PRECEPT: *Lundbeck A/S, Copenhagen, Denmark*: Misser Forrest, MD, Thomas Bisgaard, MPolSc, Erik Bardrum Nielsen, PhD, Sissel Vorstrup, MD. *Cephalon, Inc., Fraser, PA*: Heather Snyder, PhD, John Ondrasik, PhD, Lilliam Kingsbury, PhD, Steve Mulcahy, MS, Coleen Myers, BSN, Lesley Russell, MD.

Parkinson Study Group PostCEPT investigators:

Steering Committee: University of Rochester, Rochester, NY: Ira Shoulson, MD (principal investigator), Karl Kieburtz, MD, MPH, Bernard Ravina, MD, MCSE, David Oakes, PhD (chief biostatistician), Emily Flagg, BA (project coordinator), Roger Kurlan, MD; Toronto Western Hospital, University Health Network, Toronto, Ontario, Canada: Anthony Lang, MD; Rush University Medical Center, Chicago, IL: Christopher Goetz, MD; Institute for Neurodegenerative Disorders, New Haven, CT: Kenneth Marek, MD; The Parkinson’s Institute, Sunnyvale, CA: Caroline Tanner, PhD, Robin Elliot, MA; Columbia University, New York, NY: Stanley Fahn, MD.

Oversight Committee: Cephalon, Inc., Fraser, PA: Gilbert Block; Lundbeck A/S, Copenhagen, Denmark: Misser Forrest, MD; Department of Defense, Washington D.C.: Stephen Grate, DVM; The Parkinson’s Disease Foundation, New York, NY: Robin Elliot, MA; NIH/NINDS, Bethesda, MD: Diane DiEuliis, PhD, Wendy R. Galpern, MD, PhD.

PostCEPT Participating Investigators and Coordinators: Chum-Hotel Dieu/McGill Center for Studies in Aging, Montreal, Quebec, Canada: Michel Panisset, MD, Sylvain Chouinard, MD, Johanne Blais; London Health Sciences, London, Ontario, Canada: Mandar Jog, MD; Colorado Neurological Institute, Englewood CO: Dawn Miracle, BS, MS; Rush University Medical Center, Chicago, IL: Kathleen M. Shannon, MD; University of Colorado Health Sciences Center, Denver, CO: Maureen Leehey, MD, Teresa Derian, RN; University of California, Irvine, Irvine, CA and The Phillip & Carol Traub Center for Parkinson’s Disease, Eisenhower Medical Center, Rancho Mirage, CA: Neal Hermanowicz, MD, Shari Niswonger, RN; Ottawa Hospital Civic Site, Ottawa, Ontario, Canada: David Grimes, MD, RN; University of Rochester, Rochester, NY: Roger Kurlan, MD, Nancy Pearson, RN, MS; NeuroHealth Parkinson’s Disease Movement Disorders Center, Warwick, RI: Joseph Friedman, MD, Margaret Lannon, RN, MS; The Parkinson’s Institute, Sunnyvale, CA; Toronto Western Hospital, University Health Network, Toronto, Ontario, Canada; University of Minnesota/ Minnesota VA Medical Center, Minneapolis, MN: Paul Tuite, MD, Susan Rolandelli, RN; University of Iowa, Iowa City, IA: Robert Rodnitzky, MD, Judith Dobson, RN; Duke University Medical Center, Durham, NC: Burton Scott, MD, Joanne Field, BSN, RN; Oregon Health & Science University, Portland, OR: Julie Carter, RN, MN, ANP, Pamela Andrews; University of Pennsylvania, Philadelphia, PA: Andrew Siderowf, MD, Lisa Altin, BS; University of Sherbrooke, Sherbrooke, Quebec, Canada: Daniel Soucy, RN; Johns Hopkins University, Baltimore, MD: Joseph Savitt, MD, PhD, Melissa Gerstenhaber, RNC, MSN; LSU Health Science Center Shreveport, Shreveport, LA: Richard Zweig, MD, Collette Hilliard, MS; Baylor College of Medicine, Houston, TX: Joseph Jankovic, MD, Christine Hunter, RN, CCRC; Columbia University Medical Center, New York, NY: Cheryl Waters, MD, Angel Figueroa, CCRC; Northwestern University, Chicago, IL: Tanya Simuni, MD, Karen Williams; Saskatoon Dist Health Board Royal University Hospital, Saskatoon SK, Canada: Ali Rajput, MD, Marilyn Martin, BSc, ADV; University of Cincinnati/Cincinnati Children’s Hospital, Cincinnati, OH: Alberto Espay, MD, MSC; Sun Health Research Institute, Sun City, AZ: Holly Shill, MD; University of South Florida, Tampa, FL; University of Tennessee-Memphis, Memphis, TN: Ronald Pfeiffer, MD, Brenda Pfeiffer, RN, BSN; University of Virginia, Charlottesville, VA: Frederick Wooten, MD, Margaret F. Keller, RN, MS, CCRC; Albany Medical College, Albany, NY: Eric Molho, MD, Katy Regan; Boston University, Boston, MA: Marie H. Saint-Hilaire, MD, Cathi-Ann Thomas, RN, MS; Mayo Clinic Jacksonville, Jacksonville, FL: Margaret Turk, RN; Medical College of Georgia, Augusta, GA: Buff Dill, BS, ED; Milton S. Hershey Medical Center, Hershey, PA: Thyagarajan Subramanian, MD, Donna Stuppy, LPN; Institute for Neurodegenerative Disorders, New Haven, CT; UMDNJ Robert Wood Johnson Medical School, New Brunswick, NJ: Margery H. Mark, MD, Debbie Caputo, MSN, FNP-BC; University of Maryland School of Medicine, Baltimore, MD; Massachusetts General Hospital, Charleston, MA: Michael Schwarzschild, MD, PhD; University of Toledo Health Science Center, Toledo, OH: Lawrence Elmer, MD, PhD, Kathy Davis, RN Stephanie Wilson, RN, MSN, CCRC; University of Alberta, Edmonton, AB, Canada: Richard Camicioli, MD, Pamela King, BScN, RN; University of California Davis, Sacramento, CA: Lin Zhang, MD, PhD, John Bautista; University of Southern California, Los Angeles, CA; Creighton University, Omaha, NE; Paciﬁc Neuroscience Medical Group, Oxnard, CA; University of Calgary, Calgary, AB, Canada: Ranjit Ranawaya, MD; University of California San Diego, La Jolla, CA: David Song, MD, Deborah Fontaine, RNCS, MS; University of Kansas Medical Center, Kansas City, KS: Kelly Lyons, PhD, Carey Mack, RN; Beth Israel Deaconess Medical Center, Boston, MA: Daniel Tarsy, MD, Peggy Rose, RN; Brigham & Women’s Hospital, Boston, MA: Lewis Sudarsky, MD, Georgette Hage, MD; Cleveland Clinic Florida-Weston, Weston, FL: Nestor Galvez- Jimenez, MD; The Parkinson’s & Movement Disorder Institute, Fountain Valley, CA: Daniel Truong, MD; University of Chicago, Chicago, IL: Un Jung Kang, MD, Joan Young, CCRC; North Shore-LIJ Health System, Manhasset, NY: Andrew Feigin, MD, Jean Ayan, RN; Stanford University Medical Center, Stanford, CA: Helen Bronte-Stewart, MD; Beth Israel Medical Center, New York, NY: Karyn Boyar, RN, CNS, FNP; University of Arkansas for Medical Sciences, Little Rock, AR: Jana Patterson, RN.

Biostatistics and Coordination Center Staff: University of Rochester, Rochester, NY: Earl Westerlund, Lisa Lang, Tina Winebrenner, Nicole McMullen, Sandra Plumb, Cindy Casaceli, MBA.

DIGPD Study group. Steering committee: Jean-Christophe Corvol, MD, PhD (Pitié-Salpêtrière Hospital, Paris, principal investigator of DIGPD), Alexis Elbaz, MD, PhD (CESP, Villejuif, member of the steering committee), Marie Vidailhet, MD (Pitié-Salpêtrière Hospital, Paris, member of the steering committee), Alexis Brice, MD (Pitié-Salpêtrière Hospital, Paris, member of the steering committee and PI for genetic analysis) ; Statistical analyses: Alexis Elbaz, MD, PhD (CESP, Villejuif, PI for statistical analyses), Fanny Artaud, PhD (CESP, Villejuif, statistician); Principal investigators for sites (alphabetical order): Frédéric Bourdain, MD (CH Foch, Suresnes, PI for site), Jean-Philippe Brandel, MD (Fondation Rothschild, Paris, PI for site), Jean-Christophe Corvol, MD, PhD (Pitié-Salpêtrière Hospital, Paris, PI for site), Pascal Derkinderen, MD, PhD (CHU Nantes, PI for site), Franck Durif, MD (CHU Clermont-Ferrand, PI for site), Richard Levy, MD, PhD (CHU Saint-Antoine, Paris, PI for site), Fernando Pico, MD (CH Versailles, PI for site), Olivier Rascol, MD (CHU Toulouse, PI for site); Co-investigators (alphabtical order): Anne-Marie Bonnet, MD (Pitié-Salpêtrière Hospital, Paris, site investigator), Cecilia Bonnet, MD, PhD (Pitié-Salpêtrière Hospital, Paris, site investigator), Christine Brefel-Courbon, MD (CHU Toulouse, site investigator), Florence Cormier-Dequaire, MD (Pitié-Salpêtrière Hospital, Paris, site investigator), Bertrand Degos, MD, PhD (Pitié-Salpêtrière Hospital, site investigator), Bérangère Debilly, MD (CHU Clermont-Ferrand, site investigator), Alexis Elbaz, MD, PhD (Pitié-Salpêtrière Hospital, Paris, site investigator), Monique Galitsky (CHU de Toulouse, site investigator), David Grabli, MD, PhD (Pitié-Salpêtrière Hospital, Paris, site investigator), Andreas Hartmann, MD, PhD (Pitié-Salpêtrière Hospital, Paris, site investigator), Stephan Klebe, MD (Pitié-Salpêtrière Hospital, Paris, site investigator), Julia Kraemmer, MD (Pitié-Salpêtrière Hospital, site investigator), Lucette Lacomblez, MD (Pitié-Salpêtrière Hospital, Paris, site investigator), Sara Leder, MD (Pitié-Salpêtrière Hospital, Paris, site investigator), Graziella Mangone, MD, PhD (Pitié-Salpêtrière Hospital, Paris, site investigator), Louise-Laure Mariani, MD (Pitié-Salpêtrière Hospital, Paris, site investigator), Ana-Raquel Marques, MD (CHU Clermont Ferrand, site investigator), Valérie Mesnage, MD (CHU Saint Antoine, Paris, site investigator), Julia Muellner, MD (Pitié-Salpêtrière Hospital, Paris, site investigator), Fabienne Ory-Magne, MD (CHU Toulouse, site investigator), Violaine Planté-Bordeneuve, MD (Henri Mondor Hospital, Créteil, site investigator), Emmanuel Roze, MD, PhD (Pitié-Salpêtrière Hospital, Paris, site investigator), Melissa Tir, MD (CH Versailles, site investigator), Marie Vidailhet, MD (Pitié-Salpêtrière Hospital, Paris, site investigator), Hana You, MD (Pitié-Salpêtrière Hospital, Paris, site investigator); Neuropsychologists: Eve Benchetrit, MS (Pitié-Salpêtrière Hospital, Paris, neuropsychologist), Julie Socha, MS (Pitié-Salpêtrière Hospital, Paris, neuropsychologist), Fanny Pineau, MS (Pitié-Salpêtrière Hospital, Paris, neuropsychologist), Tiphaine Vidal, MS (CHU Clermont-Ferrand, neuropsychologist), Elsa Pomies (CHU de Toulouse, neuropsychologist), Virginie Bayet (CHU de Toulouse, neuropsychologist); Genetic core: Alexis Brice (Pitié-Salpêtrière Hospital, Paris, PI for genetic studies), Suzanne Lesage, PhD (INSERM, ICM, Paris, genetic analyses), Khadija Tahiri, PhD (INSERM, ICM, Paris, lab technician) Hélène Bertrand, MS (INSERM, ICM, Paris, lab technician), Graziella Mangone, MD, PhD (Pitié-Salpêtrière Hospital, Paris, genetic analyses);

Sponsor activities and clinical research assistants: Alain Mallet, PhD (Pitié-Salpêtrière Hospital, Paris, sponsor representative), Coralie Villeret (Hôpital Saint Louis, Paris, Project manager), Merry Mazmanian (Pitié-Salpêtrière Hospital, Paris, project manager), Hakima Manseur (Pitié-Salpêtrière Hospital, Paris, clinical research assistant), Mostafa Hajji (Pitié-Salpêtrière Hospital, Paris, data manager), Benjamin Le Toullec, MS (Pitié-Salpêtrière Hospital, Paris, clinical research assistant), Vanessa Brochard, PhD (Pitié-Salpêtrière Hospital, Paris, project manager), Monica Roy, MS (CHU de Nantes, clinical researh assistant), Isabelle Rieu, PhD (CHU Clermont-Ferrand, clinical research assistant), Stéphane Bernard (CHU Clermont-Ferrand, clinical research assistant), Antoine Faurie-Grepon (CHU de Toulouse, clnical research assistant).

ParkWest: Principal investigators: Guido Alves (Norwegian Centre for Movement Disorders, Stavanger University Hospital), Ole-Bjørn Tysnes (Haukeland University Hospital). Investigators and study coordinators: Karen Herlofson, Solgunn Ongre, Siri Bruun (Sørlandet Hospital Arendal); Ineke HogenEsch, Marianne Kjerandsen, Liv Kari Håland (Haugesund Hospital); Wenche Telstad, Aliaksei Labusau, Jane Kastet (Førde Hospital); Bernd Müller, Geir Olve Skeie, Charalampos Tzoulis (Haukeland University Hospital); Kenn Freddy Pedersen, Michaela Dreetz Gjerstad, Elin Bjelland Forsaa, Jodi Maple-Grødem, Johannes Lange, Veslemøy Hamre Frantzen, Anita Laugaland, Karen Simonsen, Ingvild Dalen (Stavanger University Hospital).

PICNICS: Principal investigator -Roger Barker; study team - Caroline H Williams-Gray, Jonathan R Evans, David P Breen, Gemma Cummins, Marta Camacho, Ruwani Wijeyekoon, Kirsten M Scott, Thomas Stoker, Julia C Greenland.
