## Supplementary figures and images for "Genome-wide association study of Parkinson’s disease progression biomarkers in 12 longitudinal patients’ cohorts"

### Supplemental figure 1

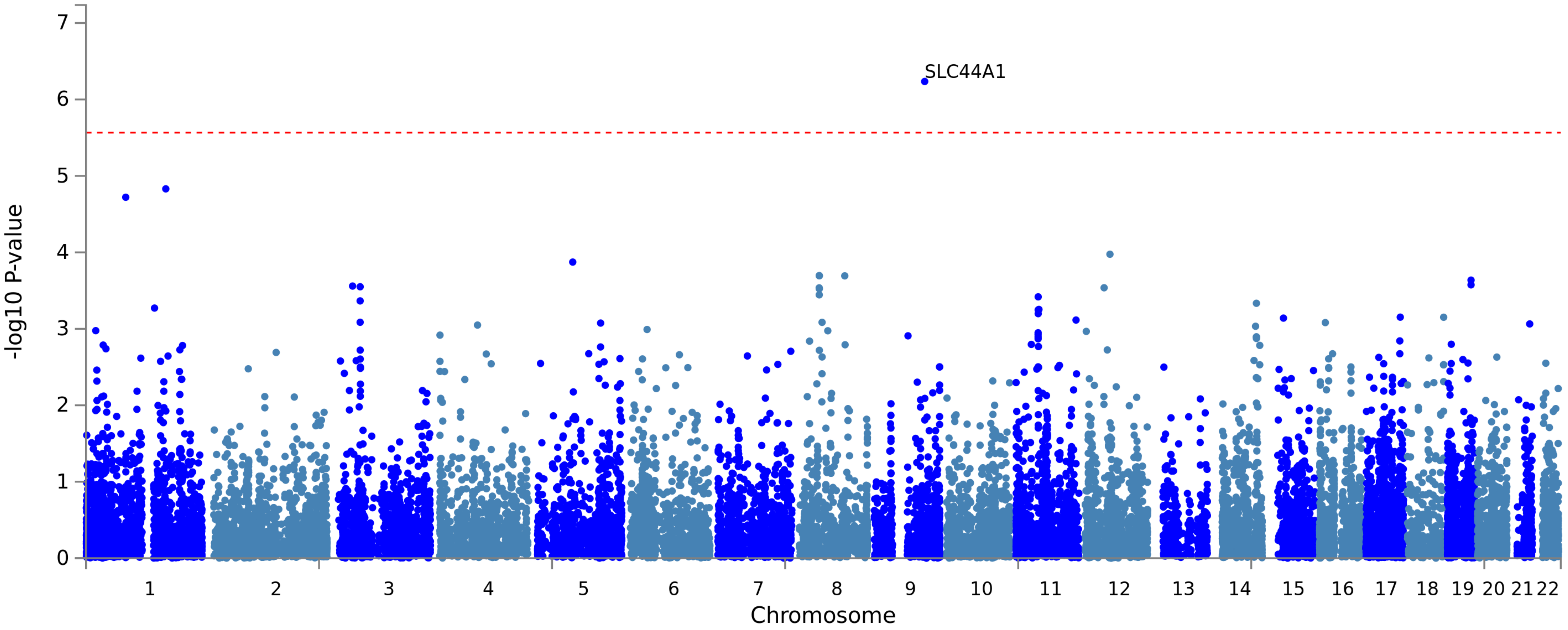

### Supplemental figure 2

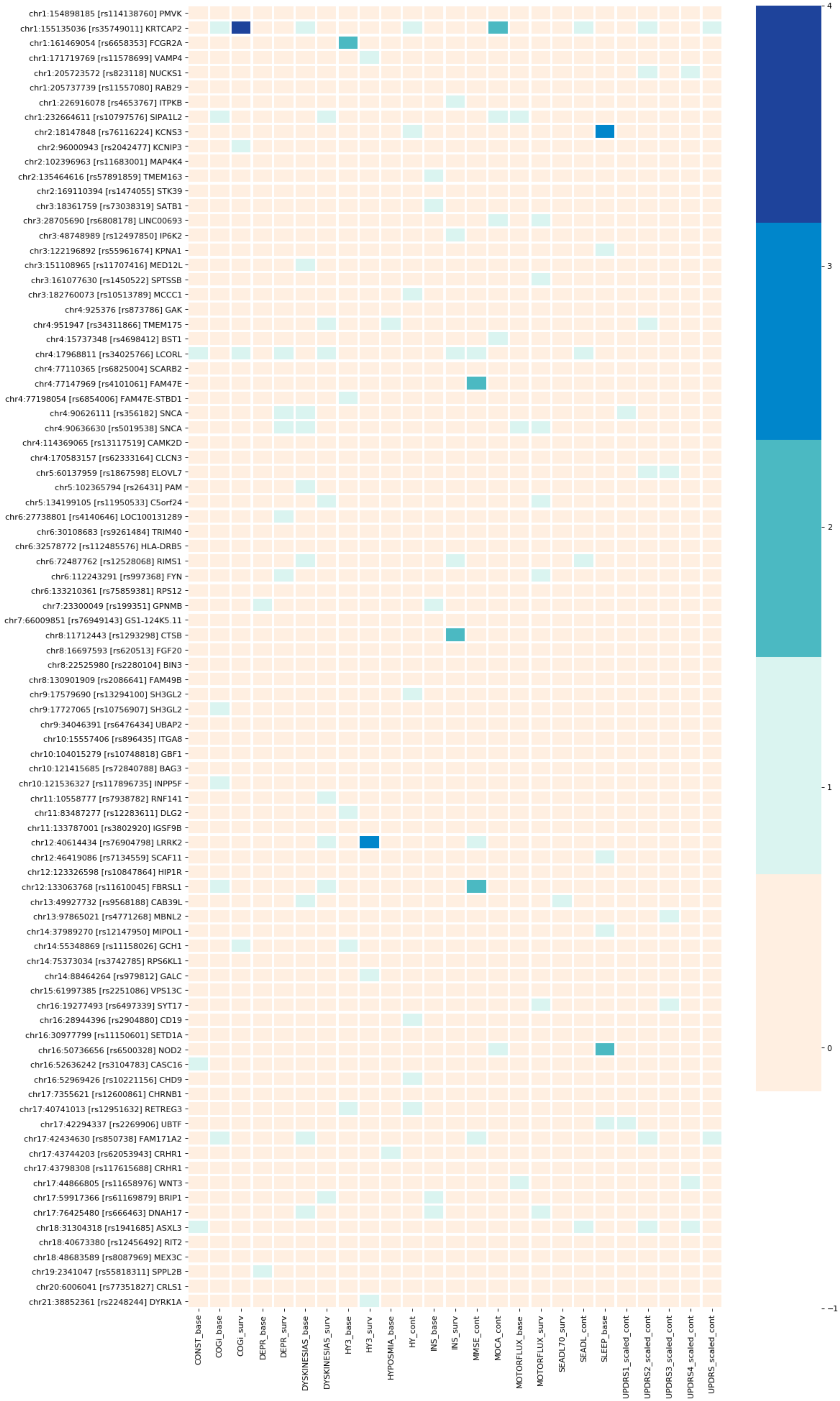
